# Non-verbal skills in auditory word processing: Implications for typical and dyslexic readers

**DOI:** 10.1101/2023.10.13.562269

**Authors:** Soufiane Jhilal, Nicola Molinaro, Anastasia Klimovich-Gray

## Abstract

Dyslexia is characterized by poor phonological skills and atypical neural responses to text and speech stimuli, while non-verbal intelligence remains unaffected. However, the impact of non-verbal intelligence on language-evoked neural responses in dyslexia is underexplored. This study examines non-verbal intelligence’s effects on neural responses to auditory words in typical and dyslexic readers. Participants completed IQ tests, reading and phonological assessments, and underwent magnetoencephalography recordings while listening to words. Event-related brain field responses at different stages of auditory word processing (100ms, 200ms, 400ms) were analyzed in relation to non-verbal IQ scores and reading performance. Higher non-verbal IQ in typical readers yielded earlier right hemisphere perceptual responses (100ms). Dyslexic readers showed no IQ-related latency effects but those with lower non-verbal IQ displayed amplified left hemisphere response amplitudes at 100ms. These findings confirm the link between non-verbal cognitive abilities and early auditory word processing in adults, highlighting distinct effects in dyslexia and controls.

## Introduction

Previous research has convincingly shown that the degree of reading impairment in dyslexia is not correlated with general intelligence skills as measured by intelligence quotient (IQ) tests (Fletcher et al., 1994; Peterson & Pennington, 2015; Shaywitz et al., 1999; Tanaka et al., 2011). According to the *core phonological deficit* view (Stanovich & Siegel, 1994), phonological impairments in dyslexia affect initial phonological stages of word processing while any worse performance for more complex linguistic tasks (e.g., text reading comprehension) are downstream consequences of these lower-level issues. Within this logic, broader domain-general intelligence, as measured by IQ tests, cannot compensate for the core phonological deficits underlying word-level reading and impact word-level processing directly. Indeed, poor readers with high and low IQ show similar phonological deficits in pseudoword reading (Stanovich & Siegel, 1994; Stanovich, 1996), phonological segmentation (Fletcher et al., 1994) and word recognition (Bruck, 1990). According to such evidence, better word-level reading and processing is a result of less severe phonological deficits and not a result of other intelligence-linked protective factors. Proponents of the *compensatory view*, however, point out that strong oral linguistic skills and domain-general intelligence factors relevant for linguistic performance, such as attention (Eklund et al., 2013), working (Holmes et al., 2015) and declarative (Ullman and Pullman, 2015) memory, as well as processing speed (Snowling, 2001) can encourage alternative compensatory mechanisms for word-level learning and reading rather than alleviate severity of the reading deficits by circumventing persistent phonological-level deficits via semantic bootstrapping (Muter & Snowling, 2009; Snowling, 2003) or memory-intensive whole word learning (Ziegler & Goswami, 2005). In this view, individual differences in some language-relevant domain-general skills could shape the cognitive strategies of word-level processing in dyslexic readers. To address this debate from a neurocognitive perspective, we used magnetoencephalography (MEG) to explore the impact of the domain-general intelligence on the neural substrates of word processing in typical and dyslexic readers. We present the first neuroimaging evidence for differential impact of non-verbal intelligence skills on the neural signatures of spoken word processing in these two groups.

### Cognitive strengths as protective factors in dyslexia

Strong verbal language skills have been proposed as protective factors, facilitating compensatory mechanisms for better word-level reading in dyslexia (Muter & Snowling, 2009; Snowling, 2003). Having superior verbal reasoning in schoolchildren with dyslexia (grades 1-9) was linked to overall better word-reading and spelling but persistently poor phonological decoding and rapid automated naming (RAN) skills (Berninger & Abbott, 2013). For instance, Van Viersen and colleagues (2014) have shown that gifted (120+ IQ) Dutch first-graders with dyslexia have outperformed average IQ dyslexic peers on verbal skills such as vocabulary and grammar yet persistently showed weaknesses in phonological tasks. These and other similar findings show a robust link between verbal skills acquired early in life and better reading outcomes in dyslexia, even if the underlying phonological deficits persist (Fletcher et al., 1994; Van Viersen et al., 2019). In the compensatory view, such effects are explained as dyslexic readers learning to rely on semantic contextual information partially attained via stronger oral linguistic skills when faced with phonological and decoding deficits (Snowling et.al., 2003, Snowling, 2008). An alternative explanation is that the gains in reading due to strong verbal skills are epiphenomenal and simply signal a less pronounced phonological deficit (Fletcher et al., 1994; Stanovich 1996) since the latter contributes to the development of both verbal and written language skills (Hulme & Snowling, 2014; Nation & Snowling, 2004; Ramus et al., 2013).

Unlike verbal language skills, non-verbal intelligence is not directly linked to development of phonological skills. Consequently, the role of non-verbal intelligence skills as potential protective factors and sources of compensatory mechanisms in dyslexia was much less explored. Several studies have made tentative links between non-verbal skills and improved performance in dyslexia. Van Viersen and colleagues (2014) showed that high IQ dyslexic first-graders had superior working and visuo-spatial memory compared to average-IQ dyslexic peers, suggesting that such domain-general skills when paired with better oral skills boost compensatory mechanisms for written text processing. Similarly, deficits in other non-verbal skills such as visuo-spatial attention (Facoetti et al., 2000; Landerl & Wimmer, 2008) and non-verbal working memory as indicated by forward and backward digit span tasks (Giofrè et al., 2016) were shown to be associated with reading difficulties in dyslexia, that is, children with higher measures in those non-verbal skills performed better in reading tasks. These studies provide tentative evidence for some non-verbal skills contributing to elevating the degree of the reading deficit in dyslexia.

Better understanding the link between non-verbal intelligence and language processing has important implications both within and outside of the dyslexia research domain. In the broader context of cognitive neuroscience of language, the question of whether non-verbal intelligence can play a compensatory role in dyslexia is directly tied to one of the key debates in the field: to what extent does language processing become domain-selective in adults, encapsulated in a dedicated pool of cognitive computation and neural resources. In line with the view that general cognition affects written and spoken language analysis at higher, more complex and resource intensive levels, numerous studies demonstrated involvement of domain-general processes in comprehension of spoken or written sentences such as attention (Hubbard & Federmeier, 2021; Ni et al., 2000; Von Kriegstein, 2003) and working memory (Emmorey et al., 2017; Gathercole & Baddeley, 2009). These domain-general processes have clear linguistic functions like the role of memory in consolidating novel words (Davis et al., 2009; Kaczer et al., 2018; Kurdziel & Spencer, 2015) and the role of attention in semantic activation and syntactic context analysis (Fischler & Bloom, 1979; Otsuka & Kawaguchi, 2007; Myachykov et al., 2005; Rogalsky & Hickok, 2008; Tomlin, 1997). This line of neuroimaging research in typical readers is complementary to the argument that greater general intelligence boosting domain-general abilities could in principle serve as a protective factor for more complex linguistic operations like sentence processing or novel word learning (Stanovich, 1996). Yet the role of the domain-general processes in word-level language operations like lexical access outside cognitively demanding task conditions (Campbell & Tyler, 2018) is a topic of debate (Klimovich-Gray et al., 2017, 2019). Therefore, based on this literature, the question of whether domain-general skills and intelligence can facilitate neural operations involved in word-level processing in dyslexia (or indeed typical readers) remains open and relevant.

Here we propose to explore the extent to which individual domain-general non-verbal cognitive abilities affect stages of auditory word processing in adults with and without dyslexic symptoms. We focus on auditory word processing since phonological deficiencies in dyslexia (Elbro et al., 1994; Martin et al., 2010; Suárez-Coalla & Cuetos, 2015; Undheim, 2009) are thought to stem from aspects of auditory speech analysis (Diaz et al., 2012; Lizarazu et al., 2021; Vandermosten et al., 2010) that result in these phonological deficits (Goswami, 2011). Furthermore, the limited evidence for domain-general intelligence having effect on linguistic skills comes from auditory processing in typically developing children (Hampton Wray & Weber-Fox, 2013; Murphy et al. 2014; Tomlin et al., 2015). If domain-general skills are linked to auditory word processing in adults with dyslexic symptoms, typical linguistically centered interventions (Christo et al., 2010) could be supplemented with non-verbal skills training which can in turn boost word-level processing and learning, leading to better remediations. To address these questions, we looked at the relationship between aspects of non-verbal (NV) IQ previously proposed as possible protective factors in dyslexia and the neural markers of different stages of single word processing. Focusing on single words avoids more complex syntactic and contextual processing contaminated by attentional and working memory demands, arguably also affected in dyslexia (Richlan, 2019; Robertson & Joanisse, 2010; Tijms, 2004).

### Effects of non-verbal skills on word processing – developmental evidence

Another important avenue to consider are developmental studies showing that effects of domain-general abilities on language-related skills are present in young children (Hampton Wray & Weber-Fox, 2013; Murphy et al. 2014; Tomlin et al., 2015) in basic auditory paradigms (Baltes & Lindenberger, 1997; Hampton Wray & Weber-Fox, 2013; Lindenberger & Baltes, 1994; Scott Acton & Schroeder, 2001). One of the hypothesized reasons for this is that the immature neural circuits of children do not yet differentiate between different cognitive domains (Booth et al., 2001).

Evidence from behavioral studies in children has shown that NV IQ is related to auditory processing and lexical access. NV IQ was positively correlated with measures assessing auditory processing performance in children (7–12 years old), namely a frequency pattern, a dichotic digits test, and a listening in spatialized noise-sentences test (Tomlin et al., 2015). Their NV IQ scores were also a significant predictor of listening ability which suggests that NV IQ plays a role in facilitating auditory processing of language at least in children. Similarly, small but positive correlations were found between general intelligence and measures of pitch discrimination in teens and adults (Scott Acton & Schroeder, 2001). Intelligence was also found to be related to auditory acuity (Baltes & Lindenberger, 1997; Lindenberger & Baltes, 1994). Conversely, others failed to find any effects for NV IQ on auditory processing or lexical access. Abrigo (2012) assessed lexical processing speed in children and adults by measuring the latency of eye fixation to pictures of target nouns presented auditorily. In both children and adults neither the full-scale IQ score nor the verbal and non-verbal indices of the Wechsler IQ test showed significant effects on the lexical processing speed. Murphy et al. (2014) also found no effect of NV IQ on auditory processing in children when examining correlations between the RAVEN test of Progressive Matrices and auditory temporal tests (Frequency Pattern and Gaps in Noise).

The existing behavioral evidence is mixed and inconclusive. Furthermore, most studies do not explore which stages of processing were specifically linked to NV IQ abilities. We can address this by examining the underlying time-resolved brain activity with electro- or magnetoencephalographic data using event related potentials (Malins et al., 2013), where dissociable evoked components have been associated with different stages of word processing, (Kaan, 2007). The early 100ms response is elicited by the onset of perception of the auditory stimuli (Näätänen & Picton, 1987), the 200ms component indexes early phonological and orthographic activation (Kong et al., 2010) while the later component peaking at 400ms reflects stimulus lexico-semantic evaluation and classification (Sur & Sinha, 2009). Hence, in this study we asked if and how auditory word-processing evoked responses are modulated by non-verbal cognition indices.

Several EEG studies exploring these questions support behavioral evidence linking domain-general NV IQ abilities to early stages of auditory word processing in children. Using EEG, An et al. (2020) investigated children’s (3–8 years old) responses to human voice and found IQ to be correlated with the P100 response amplitude and gamma oscillatory power changes, consistent with early behavioral work (Baltes & Lindenberger, 1997; Lindenberger & Baltes, 1994; Tomlin et al., 2015). Effects of NV IQ in children (7–9 years old) were also found on the N400 and P600 components in sentences with context effects (Hampton Wray & Weber-Fox, 2013). Given that there are no corresponding studies in adults, it is possible that domain-general non-verbal abilities only have an impact on the neural signatures of auditory word processing in children. Unlike adults who have a matured cognitive system, children are thought to rely on general cognition until they can develop specialized systems for different tasks (Booth et al., 2001). Effects of general cognition on language processing are almost never explored in adults since almost all language related studies tend to control for IQ as a covariate of no interest.

### Significance of current work for dyslexia research

Currently there is no work exploring the above questions in adult populations with language and speech processing impairments. Understanding the relationship between domain-general and linguistic skills in these populations is critical as such links can be explored in remediation settings. Dyslexic readers are a perfect target group to address these questions since they exhibit poor reading and phonological skills despite largely normal domain-general NV IQ abilities (Peterson & Pennington, 2015; Ram us, 2003; Kovelman et al., 2011; MacSweeney et al., 2009; Snowling, 2008). Recent work has highlighted that both reading and precursory phonological deficits can stem from early and persistent problems in accessing and retrieving low-level acoustic inputs (Boets et al. 2013; Boets, 2014; Ramus, 2014; Vandermosten et al., 2013; Diaz et al., 2012; Lizarazu et al., 2021), including difficulties in perceptual sampling of the auditory input (Goswami, 2011) that can be measured in infancy (Kalashnikova et al. 2018). If domain-general abilities in this group are linked to auditory word processing measures, this could open avenues for supplementary domain-general skills interventions from the earliest stages of speech acquisition.

In line with this, EEG studies have found that subjects with dyslexia have reduced auditory evoked N100 amplitudes (Bonte & Blomert, 2004) and prolonged latencies (Molfese, 2000) as well as P300 component latency and amplitude abnormalities (Papagiannopoulou & Lagopoulos, 2016) when compared to controls. Papageorgiou et al. (2009) also reported significantly reduced N100 amplitudes for dyslexic children compared to their sibling controls for pure tone perception. Critically, memory performance in the control group was positively correlated with N100 amplitude but no similar effect was observed for the dyslexic group. This could suggest that at least for children with dyslexia domain-general skills have a weaker link with auditory processes, compared to controls. However, since this study used pure tones, it is not possible to say whether the same holds true in the context of auditory language processing.

### Current study

The current study explores the relationship between NV IQ, and the latency and amplitude of the 100ms, 200ms and 400ms event-related field (ERF) responses (collected using MEG) elicited by auditory word processing in typical and dyslexic readers. These three ERF components allow us to explore different levels of auditory word processing starting from the early auditory (M100), to phonological (M200) and lexico-semantic processing stages (M400). Many studies investigating NV IQ using the WAIS IQ test used the Performance IQ Index (PIQ) from the WAIS-III version (Wechsler, 1997) as a measurement of NV IQ (Bishop et al., 2014; Kimura & Harshman, 1984; McLaurin et al.; 1973). The WAIS-III version was composed a Performance IQ Index and a Verbal IQ (VIQ) index; however, these two performance/verbal indices were dropped in subsequent versions of the WAIS IQ test and replaced by four index scores representing more narrow domains of cognitive function which are: the Verbal Comprehension Index (VCI), the Perceptual Reasoning Index (PRI), the Working Memory Index (WMI), and the Processing Speed Index (PSI). VCI measures the ability to understand, learn and retain verbal information and to use language to solve novel problems while WMI measures the ability to hold verbal information in short-term memory and to manipulate that information. Both indices are not appropriate measurements of NV IQ in the context of our experimental questions since they measure the individual’s ability to process verbal information. As for PRI, it measures the ability to understand visual information and to solve novel abstract visual problems by scoring the subject’s performance on block design, matrix reasoning and visual puzzle tasks. These tasks require the individual to understand abstract and complex visual information which requires intensive resources of visual attention, a capacity shown to be deficient in dyslexia (Collis et al., 2013; Van der Lubbe et al., 2019; Zhao et al., 2018), and thus not a proper measure of unaffected non-verbal skills. On the other hand, processing speed, which is measured by PSI in the WAIS-IV IQ test (Wechsler, 2008) using symbol search (looking at two target symbols and then examining a group of symbols to see if the target symbols are repeated) and coding (recording associations between different symbols and numbers within time limits) tasks, is not typically impaired in dyslexia (Snowling, 2001), and thus would be a proper measure of unaffected non-verbal cognitive resources. Accordingly, and to isolate discrete non-verbal cognitive function from verbal and perceptual domains, NV IQ will be assessed with the Processing Speed Index (PSI) score of the WAIS-IV IQ test (Wechsler, 2008).

Based on previous work done in children (Hampton Wray & Weber-Fox, 2013; Murphy et al. 2014; Tomlin et al., 2015), we hypothesize that greater NV IQ abilities (as measured by PSI) will result in earlier latencies in adults, especially for early sensory processes captured by the M100 component. If later stages of lexical access are more independent from non-verbal domain-general resources, we expect weaker or no modulation of the M200 and M400 component by the NV IQ skills. Furthermore, we hypothesize that NV IQ will have a different relationship with auditory word processing in dyslexic readers compared to controls. Here, two opposite scenarios are possible. First, based on limited research in children (Papageorgiou et al. 2009) we can expect weaker effects of NV IQ on auditory processing in dyslexic adults. This could be because dyslexic readers have deficits in mechanisms involved in early selective attention, which could limit their ability to use non-verbal cognitive resources for faster auditory processing. Conversely, stronger effects of NV IQ on auditory word analysis in dyslexic participants are also possible if, unlike children, dyslexic adults can use domain-general resources in a compensatory manner. This could indicate that as neural circuits mature and all specialized systems are established, the brain learns to adapt and counteract deficits in auditory word processing by leveraging non-verbal domain-general skills.

## Methods

### Participants

We collected data from 45 Spanish native speakers with normal or corrected-to-normal vision, normal hearing, and no history of neurological or psychiatric disorders. Due to incomplete behavioral and/or MEG protocols or noisy signal, data from 3 participants were excluded. This experiment was part of a larger experiment approved by the Basque Center on Cognition Brain and Language (BCBL) ethical committee (see Klimovich-Gray et all., 2021). All participants signed an informed consent form and were paid for their time. In order to identify dyslexic participants, subjects had to meet all the following criteria: a) normal IQ (full IQ superior to 80 on WAIS-IV IQ test); b) previous formal diagnosis of dyslexia (where possible) or self-reported reading difficulties; c) significantly impaired scores in reading skills (see section 2.3. Behavioral tests for details). According to these criteria, we identified 12 dyslexic participants (4 males; 8 females) who were matched to the 12 controls (4 males; 8 females) on both age are relevant IQ parameters for the subsequent analyses of this study. This sample size of dyslexic participants is comparable to several previously published neuroimaging studies (Hämäläinen et al., 2012 - 11 dyslexic adults; Klimovich-Gray et al., 2023 - 14 dyslexic adults; Molinaro et al., 2016 - 10 dyslexic adults; Power et al., 2016 - 12 dyslexic children), and was previously shown to be sufficient to demonstrate replicable differences between control and dyslexic groups. Thus, this sample size was deemed sufficient for the planned analyses.

### Stimuli

The stimuli consisted of 80 single Spanish words. All words were frequent nouns (word frequency at least 8 per million) with high familiarity, imageability and concreteness ratings (min 5 out of 7), 2-3 syllables in length, taken from the ESPAL database (Duchon et al. 2013). The words we recorded by a female Spanish native speaker on a digital recorder (Marantz PMD) with sampling rate of 44kHz. All audio files were normalized with respect to their loudness using the ffmpeg-normalize software (https://github.com/slhck/ffmpeg-normalize) and EBU R128 normalization method to ensure that all files have the same perceived audio volume (−15.3 dB LUFS).

### Behavioral tests

All participants were administered on separate days a battery of behavioral tests which included: WAIS-IV IQ test, PROLEC-SE-R (Spanish reading test for adolescents), RAN and several additional tests of phonological abilities (phonological deletion, phonological short-term memory, not discussed in the present analysis). For the current set of questions and analysis, we only used scores for WAIS-IV IQ test as main independent measures and PROLEC-SE-R (Cuetos et al., 2016).

Full Scale IQ, Verbal Comprehension Index (VCI), Perceptual Reasoning Index (PRI), Working Memory Index (WMI), and Processing Speed Index (PSI). VCI, PRI, WMI, and PSI scores were recorded for all participants. PSI was used as the measurement of NV IQ while VCI was used as the measurement of verbal IQ to be controlled for during the analysis. These scores are provided in Table 1.

**Table. 1.** Means and standard deviations (SD) for age, Full Scale IQ, VCI, PRI, WMI, PSI, PROLEC-SE-R main score and Pseudo-words subtest. Measures where dyslexic participants are significantly different from controls are highlighted in bold.

|  | All participants | Controls | Dyslexic |
| --- | --- | --- | --- |
| Age | 30.02 (9.73) | 28.66 (9.31) | 30.41 (9.64) |
| Full Scale IQ | 103.61 (13.09) | **109.25 (11.52)** | **94.83 (11.44)** |
| VCI | 100.24 (16.56) | 104.66 (14.60) | 94.16 (18.16) |
| PRI | 106.68 (13.40) | 108.25 (14.63) | 102.83 (11.64) |
| WMI | 98.68 (15.62) | **109.08 (13.10)** | **84.75 (6.82)** |
| PSI | 105.87 (14.87) | 107.66 (15.55) | 102.08 (12.80) |
| PROLEC-SE-R | 97.76 (17.15) | **109.33 (11.81)** | **77.90 (9.09)** |
| Pseudo-words | 97.36 (29.00) | **115.50 (23.45)** | **66.09 (16.14)** |

Participants were administered the PROLEC-SE-R with all its subtests. The PROLEC-SE-R main score and the Pseudo-words subtest score were used as measurements of reading skills to select those with persistent dyslexic symptoms (scores displayed on Table 1). Some adults learn strategies to compensate for their initial reading difficulties, and do not obtain significantly low scores on PROLEC-SE-R main score. However, difficulties with reading pseudo-words were found to persist into adulthood (Hanley, 1997; Parrila et al., 2007; Pennington et al., 1990; Snowling et al., 2007). Therefore, significantly low scores at either the PROLEC-SE-R main score or the Pseudo-words subtest score were used as criteria to identify those with persistent dyslexic symptoms. The individual performance of each dyslexic participant was compared to the average performance of the control group using one-tailed modified t tests (Crawford & Howel, 1998) adapted to assess deficits at the individual level when the normative sample’s size is small. As such, participants with a score below or equal to −1.82 (cutoff value to attain significance) were defined as dyslexic. According to these criteria, 12 dyslexic subjects were identified. A group of 12 controls that matched the dyslexic participants in age (t= −0.452, p= 0.656), PSI (t= 0.960, p= 0.348) and PRI (t= 1.004, p= 0.327) was picked from the main sample. They were not matched on WMI because dyslexic readers are shown to display working memory deficiencies compared to controls (Beneventi et al., 2010; Fostick & Revah, 2018; Palmer, 2000; Smith-Spark & Fisk, 2007). They were also not matched on VCI since dyslexic participants are expected to be worse than controls in verbal skills, and therefore the full-scale IQ score is also expected to be lower for dyslexic participants since it encompasses VCI and WMI.

To evaluate the effect of NV IQ on single word processing, participants were divided into groups based on their scores in PSI using a median split, into a higher NV IQ and a lower NV IQ group. The higher and lower NV IQ groups in each condition (controls or dyslexic readers) were matched in age and all IQ indices (VCI, WMI and PRI) but PSI in order to control for any confounding effects. As can be seen in Table 2, controls only showed significant differences between lower NV IQ and higher NV IQ groups in PSI (t=4.669, p<.001). As for dyslexic readers, the lower NV IQ and higher NV IQ groups differed only in PSI (t=4.353, p=0.001) and Full-Scale IQ (t=3.095, p=0.011) which was driven by the difference in PSI. There were no significant differences between lower NV IQ and higher NV IQ groups in in any other behavioral measure. In addition to that, we also made sure that controls and dyslexic participants were still matched in age, PSI and PRI according to their NV IQ splits. Accordingly, there were no significant differences in age (t= 0.578, p= 0.576), PSI (t= 1.628, p= 0.134) and PRI (t= −0.688, p= 0.507) between the dyslexic higher NV IQ group and the control higher NV IQ group. There were also no significant differences in age (t=−1.042, p= 0.322), PSI (t= 0.639, p= 0.537) and PRI (t= 2.098, p= 0.062) between the dyslexic lower NV IQ group and the control lower NV IQ group.

**Table. 2.** Means, standard deviations (SD) and t tests for age, Full Scale IQ, VCI, PRI, WMI, PSI, PROLEC-SE-R main score and Pseudo-words subtest for lower NV IQ and higher NV IQ groups. Significant t tests are highlighted in bold.

| **Matched controls** |  |  |  |
| --- | --- | --- | --- |
|  | **Higher NV IQ** | **Lower NV IQ** | **t tests** |
| **Age** | 30.00 (8.46) | 27.33 (10.72) | t=0.478, p=0.643 |
| **Full Scale IQ** | 109.16 (10.30) | 109.33 (13.63) | t=-0.024, p=0.981 |
| **VCI** | 103.00 (12.31) | 106.33 (17.63) | t=-0.380, p=0.712 |
| **PRI** | 101.50 (13.80) | 115.00 (13.06) | t=-1.739, p=0.113 |
| **WMI** | 107.16 (14.42) | 101.00 (12.68) | t=-0.489, p=0.636 |
| **PSI** | 120.00 (7.43) | 95.33 (10.59) | **t=4.669, p<.001** |
| **PROLEC-SE-R** | 114.33 (11.62) | 104.33 (10.57) | t=1.559, p=0.150 |
| **Pseudo-words** | 121.83 (30.13) | 109.16 (14.33) | t=0.930, p=0.374 |
| **Matched dyslexic readers** |  |  |  |
|  | **Higher NV IQ** | **Lower NV IQ** | **t tests** |
| **Age** | 27.16 (8.51) | 33.66 (10.32) | t=-1.189, p=0.262 |
| **Full Scale IQ** | 102.50 (9.69) | 87.16 (7.30) | **t=3.095, p=0.011** |
| **VCI** | 102.83 (21.74) | 85.50 (10.42) | t=1.760, p=0.109 |
| **PRI** | 106.33 (10.25) | 99.33 (12.80) | t=1.046, p=0.320 |
| **WMI** | 88.50 (4.41) | 81.00 (7.01) | t=2.216, p=0.051 |
| **PSI** | 112.00 (9.46) | 92.16 (5.91) | **t=4.353, p=0.001** |
| **PROLEC-SE-R** | 84.83 (4.19) | 69.60 (5.22) | **t=5.418, p<.001** |
| **Pseudo-words** | 71.66 (11.63) | 59.40 (19.47) | t=1.297, p=0.227 |

### Procedure and MEG data acquisition

Auditory evoked magnetic fields were recorded in a magnetically shielded room with a whole-scalp MEG system (Elekta Neuromag, Helsinki, Finland) located at the BCBL in Donostia-San Sebastián, with a 0.03 – 330 Hz bandpass filter, and a sampling rate of 1 kHz. The system is equipped with 306 uniformly distributed channels - 102 magnetometer and 204 gradiometers. Four Head Position Indicator (HPI) coils were used to monitor subjects’ head position. The location of each coil relative to the anatomical fiducials (nasion, left and right preauricular points) was defined with a 3D digitizer (Fastrak Polhemus, Colchester, VA, USA). Eye movements were monitored with two pairs of electrodes in a bipolar montage placed on the external sides of each eye (horizontal electrooculography (EOG)) and above and below the right eye (vertical EOG). Cardiac rhythm was monitored using two electrodes, placed on the right side of the participants’ abdomen and below the left clavicle.

The data for the current analysis was taken from a larger experiment, where each participant performed 3 blocks of data acquisition. In the first block, resting state MEG activity was recorded for 5 minutes with the participants’ looking at a blank screen with their eyes open. The second block consisted of participants listening to sentences. The third block used in this analysis (final block) consisted of passive listening to 80 target words with no task. All auditory stimuli were delivered with a random inter-stimulus interval (ISI) (from 1 to 2.5 seconds) via non-magnetic plastic tubes.

### Pre-Processing

MEG data were processed using Minimum Norm Estimation (MNE) Python pipeline (Gramfort et al., 2013, version 0.21). First, we used the Signal Space-Separation (SSS) method (Taulu et al., 2005) implemented in MaxFilter 2.2 in order to remove external magnetic noise from the MEG recordings. Bad channels detected during the acquisition, were cross-checked manually and substituted using MNE python’s field interpolation method for the MEG channels. This method uses field mapping algorithms and generates minimum-norm projection of the channel signals to the sphere and back to estimate responses in the bad channels. Only good channels are projected and used for interpolation. All sensor space subject-specific data was transformed to the second block of that subject’s data. The data were further high-pass filtered at 0.1 Hz and lowpass filtered at 40 Hz. EOG and ECG artifacts were detected and subtracted from the MEG data using Independent Component Analysis (ICA) implemented in MNE Python based on high correlations with the data from the vertical and horizontal EOG electrodes and the ECG electrodes. The ICA decomposition was performed using the FastICA algorithm (Hyvärinen & Oja, 2000) and 3.17 components (SD=0.66) were removed per participant on average. Subsequently, data were segmented into epochs from −100 to 700ms aligned to the onset of the target word. Noisy epochs with high sensor amplitude values (exceeding 4000 ft in magnetometers or 4000 ft/cm in gradiometers) were excluded. On average, 76.11 epochs (SD=10.52) for each participant were included after rejections. Finally, baseline correction was also applied to the epochs data using the 100ms window prior to stimulus presentation.

### Data analysis

After Maxfilter, magnetometers and gradiometers encode highly correlated information (Garcés et al., 2017). Therefore, we only used gradiometers as they are typically less noisy. As there are two gradient measurements per sensor location, the values of each gradiometer pair were calculated as the root-mean-square (RMS) per pair. With a longitudinal split at the middle, sensors at each half were associated to their respective hemispheres. All gradiometer pairs belonging to each hemisphere were used with no specific regions of interest. Subsequently, ERF waveforms were calculated using global field power (GFP) per hemisphere, and for each subject. GFP is the standard deviation across all channels as a function of time and is used to quantify the instantaneous global activity across the spatial potential field over the scalp. The result of this analysis is a waveform that represents the temporal changes in the GFP.

We used the GFP signal averaged over all participants to determine the segments between which the 100ms, 200ms and 400ms ERF responses were bound. The latencies of the M100, M200 and M400 components were measured using the fractional area method (Hansen & Hillyard, 1980; Luck, 2005) in segments between 75 and 155 ms (M100), 155 and 270 ms (M200) and between 270 and 500 ms (M400) post stimulus. This technique defines the latency of the component as the first time point at which 50% of the total area of the component has been reached. Fractional area latency is more efficient and noise resistant than peak latency (Clayson et al., 2013). The amplitude of each component was measured using the mean amplitude approach which averages the ERF amplitudes between two fixed time points (e.g., onset and offset of an ERF component). This approach was found to be the most robust amplitude extraction method against increases in background noise (Clayson et al., 2013; Luck; 2005).

### Statistical analysis

Repeated measures ANOVAs by hemisphere were conducted on the matched sample of controls and dyslexic readers for each ERF component’s amplitude and latency, with the NV IQ subgroups and controls/dyslexic grouping as between subject factors in order to reveal any NV IQ effects specific to only one of the two subpopulations. The Holm–Bonferroni (Holm, 1979) method was used to correct P-values for multiple comparisons (across multiple ERFs tested). Supplementary hierarchical regressions were also used to verify linear relationships between NV IQ and latencies or amplitudes with Age and VCI already controlled for. Correlation analyses were also used to further examine any significant effects revealed by the ANOVAs revealed linear relationships between examined factors. All assumptions were verified for each component as the observations are independent and the variables follow a normal distribution.

## Results

Across all ANOVAs conducted, the only significant effects linked to the group differences groups were the following: in the controls’ subgroup M100 component’s latency was linked to NV IQ levels, while the dyslexic subgroup NV IQ levels affected M100 component’s amplitude - see Fig. 1. The amplitudes’ ANOVA plots are displayed on Fig. 2. We break down all significant effects below.

**Figure 1:**
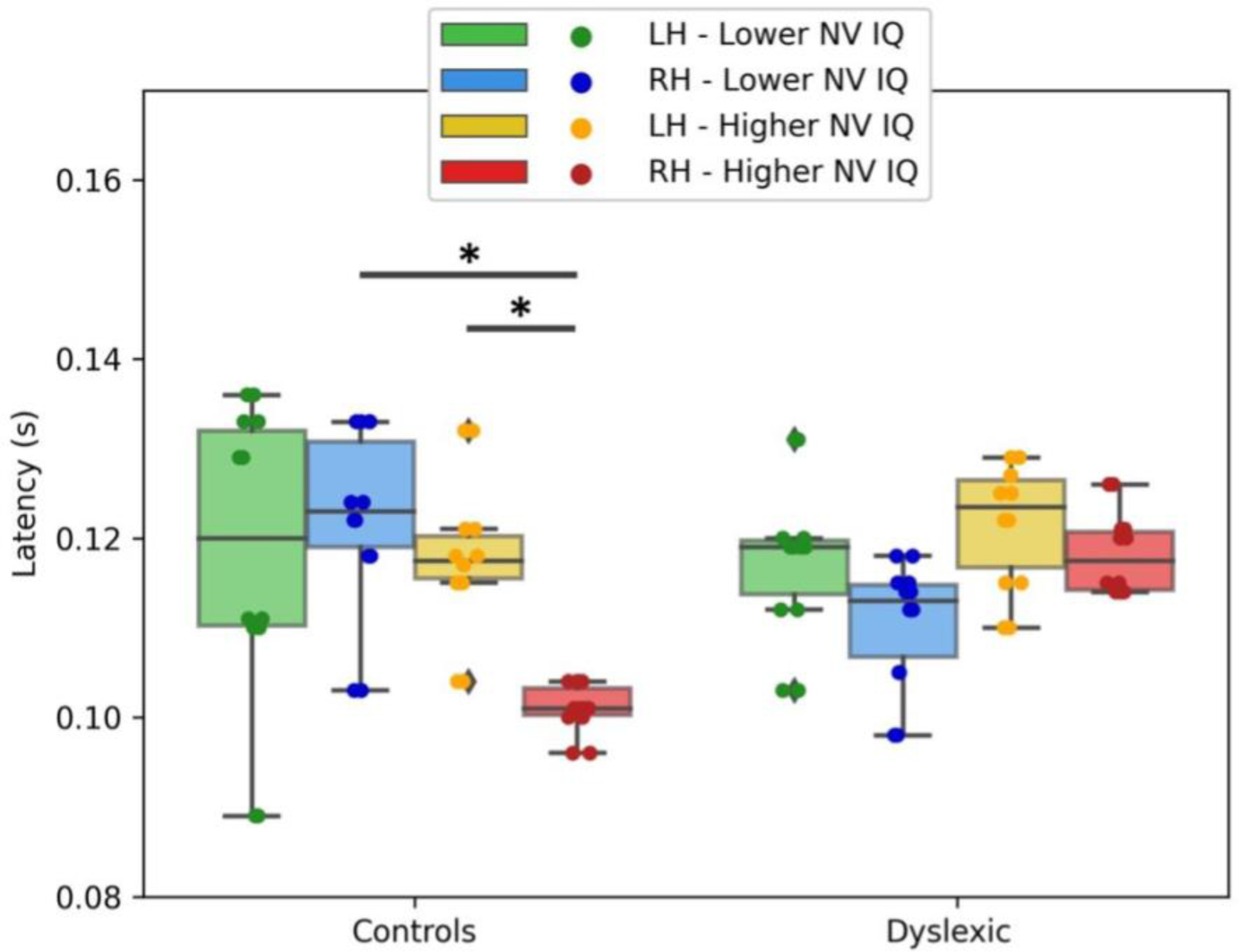
Fractional latencies (with standard errors) of the M100 component split by controls/dyslexic, hemispheres, and NV IQ subgroups. Asterisks (*) indicate significant effects.

**Figure 2:**
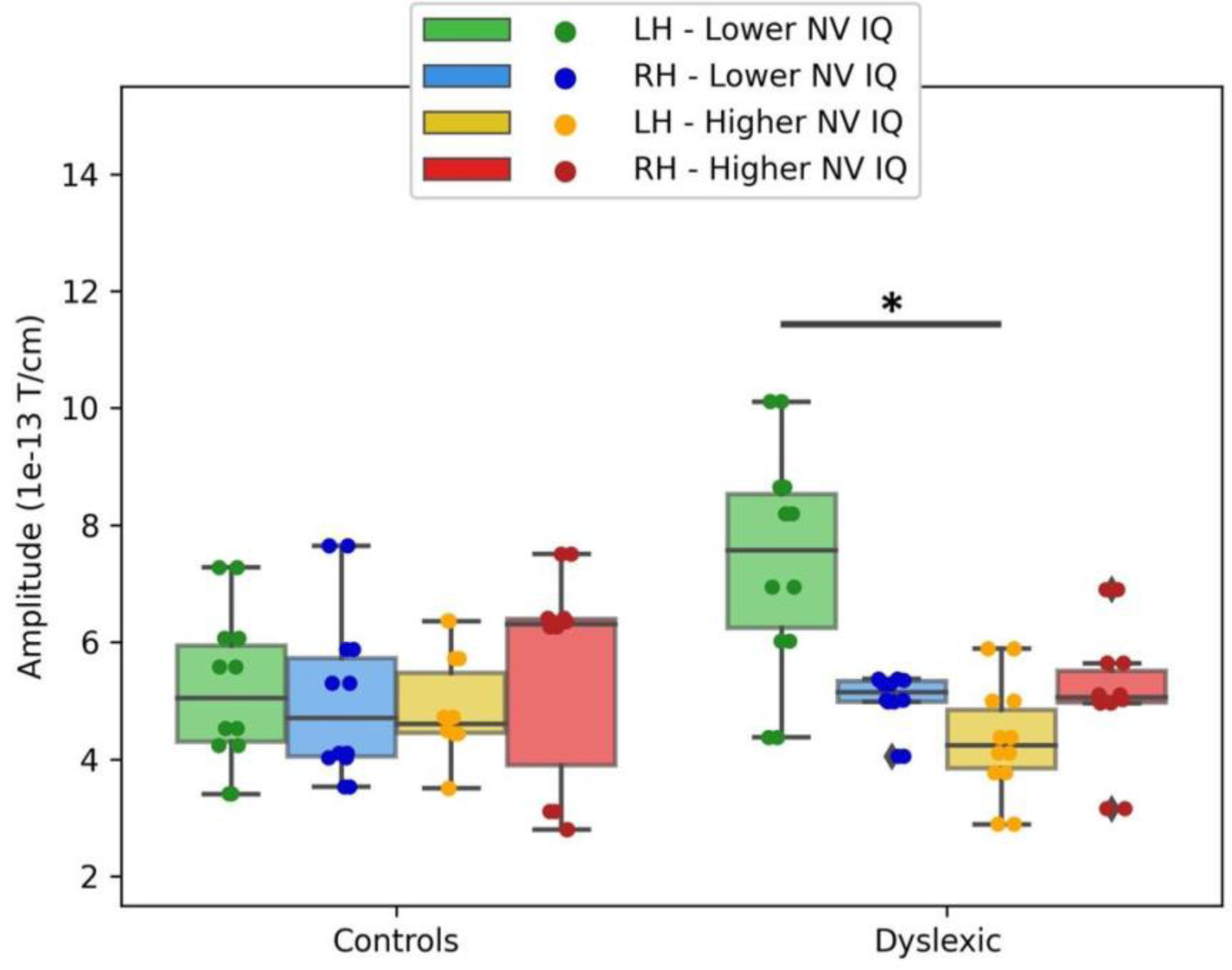
Mean amplitudes (with standard errors) of the M100 component split by controls/dyslexic, hemispheres, and NV IQ subgroups. Asterisks (*) indicate significant effects.

### M100

The analysis of variance for the M100 component showed significant effects on latency for controls and on amplitude for dyslexic readers. Regarding latency, three significant effects appeared. The effect of Hemispheres, F(1, 20) = 6.323, p = 0.021, showed that the RH was overall faster than the LH. However, the 3-way interaction Hemispheres*NV IQ*Controls/Dyslexic, F(1, 20) = 7.692, p = 0.012, showed that only for controls with higher NV IQ, the RH was significantly faster than the LH, t = −3.735, Holm-Bonferroni corrected p = 0.034. Higher NV IQ controls’ RH was also significantly faster than lower NV IQ controls’ RH, t = −3.760, Holm-Bonferroni corrected p = 0.017. These effects are clearly pronounced on both the topographic maps (Fig. 3.a) and the ERF waveforms (Fig. 3.c) of Controls while no such effects appear on the topographic maps (Fig. 3.b) and the ERF waveforms (Fig. 3.d) of dyslexic readers. Finally, the significant interaction of NV IQ*Controls/Dyslexic, F(1, 20) = 6.450, p = 0.020, was driven by the 3-way interaction above. The main effects of Controls/Dyslexic, F(1, 20) = 0.403, p = 0.533, and NV IQ, F(1, 20) = 0.506, p = 0.485, and the interactions of Hemispheres*Controls/Dyslexic, F(1, 20) = 0.088, p = 0.770, and Hemispheres*NV IQ, F(1, 20) = 3.557, p = 0.074, were all not significant.

**Figure 3:**
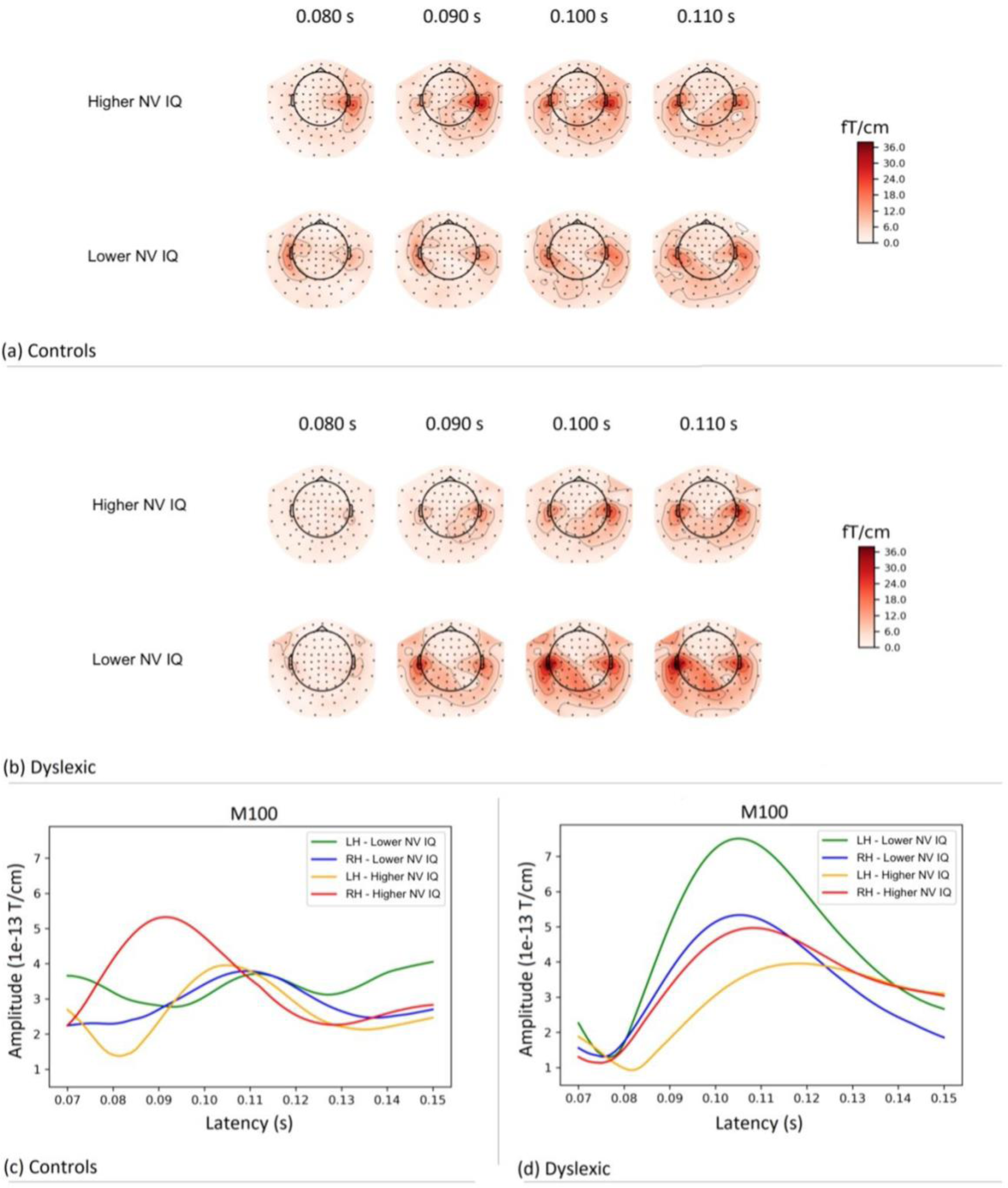
Gradiometer topographic maps and grand average waveforms showing the different effects of NV IQ on the M100 component in controls and dyslexic readers. (a) Gradiometer topographic maps of the M100 component across 4 time points for each NV IQ subgroup in controls. (b) Gradiometer topographic maps of the M100 component across 4 time points for each NV IQ subgroup in dyslexia. (c) Grand average M100 waveform per hemisphere and per NV IQ subgroup in controls. (d) Grand average M100 waveform per hemisphere and per NV IQ subgroup in dyslexia group.

In addition to the ANOVA results, to graphically illustrate the dependencies within the data, below we plot the correlation of NV IQ and the M100 latency in the right hemisphere separately for controls and dyslexic subjects (Fig. 4). A robust correlation was evident for controls, r = −0.876, p < 0.001, but not for dyslexic readers, r = 0.396, p = 0.202. This effect of NV IQ on the RH M100 latency was robust even after accounting for the effects of age, verbal IQ (as measured with VCI) and the other IQ components (WMI & PRI) in a hierarchical regression (see supplementary).

**Figure 4:**
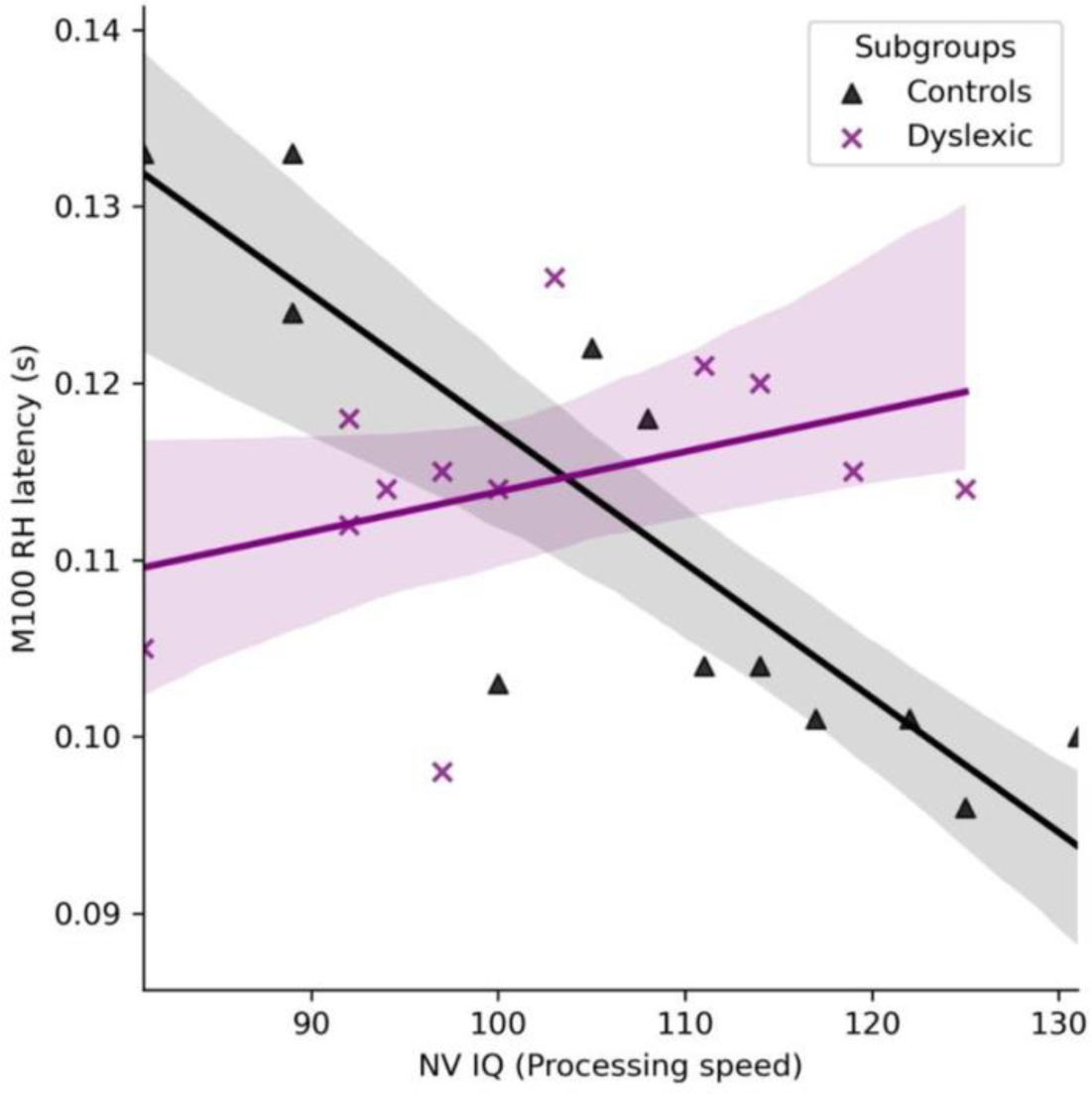
Scatter plot and correlations between NV IQ and the M100 right hemisphere (RH) latency, split by the controls/dyslexic grouping.

As for the M100 amplitudes, the main effects of Hemispheres, F(1, 20) = 0.560, p = 0.463, Controls/Dyslexic, F(1, 20) = 0.576, p = 0.457, and NV IQ, F(1, 20) = 2.808, p = 0.109, were not significant. The interactions of Hemispheres*Controls/Dyslexic, F(1, 20) = 1.714, p = 0.205, Controls/Dyslexic*NV IQ, F(1, 20) = 2.896, p = 0.106, and the 3-way interaction Hemispheres*Controls/Dyslexic*NV IQ, F(1, 20) = 2.713, p = 0.115, were all not significant. The Hemispheres*NV IQ interaction, F(1, 20) = 6.120, p = 0.022, was the only significant effect. Post hoc analysis of this interaction revealed a significant difference between the M100 left hemisphere amplitude of the higher NV IQ and the lower NV IQ subgroups, t = −2.895, Holm-Bonferroni corrected p = 0.037. Further analysis showed that this interaction was only driven by the dyslexia group - the M100 amplitude in the left hemisphere was higher for dyslexic readers with lower NV IQ compared to dyslexic readers with higher NV IQ, t = −3.716, Holm-Bonferroni corrected p = 0.018. This effect can be observed on the topographic maps (Fig. 3.b) and the M100 ERF waveforms (Fig. 3.d) where amplitudes on the left hemisphere are higher for the lower NV IQ subgroup compared to the higher NV IQ subgroup in dyslexic readers. However, even though there is a noticeable trend on the scatter plot distribution, (see Fig. 5.), there is no evidence that there is a linear relationship between NV IQ and LH M100 amplitude in dyslexic readers – see the correlation plot on the Fig. 5 below, r = −0.558, p = 0.059. A hierarchical regression between NV IQ and the LH M100 amplitudes in the dyslexic group, after accounting for the effects of age, verbal IQ (as measured with VCI) and the other IQ components (WMI & PRI), was also not significant.

**Figure 5:**
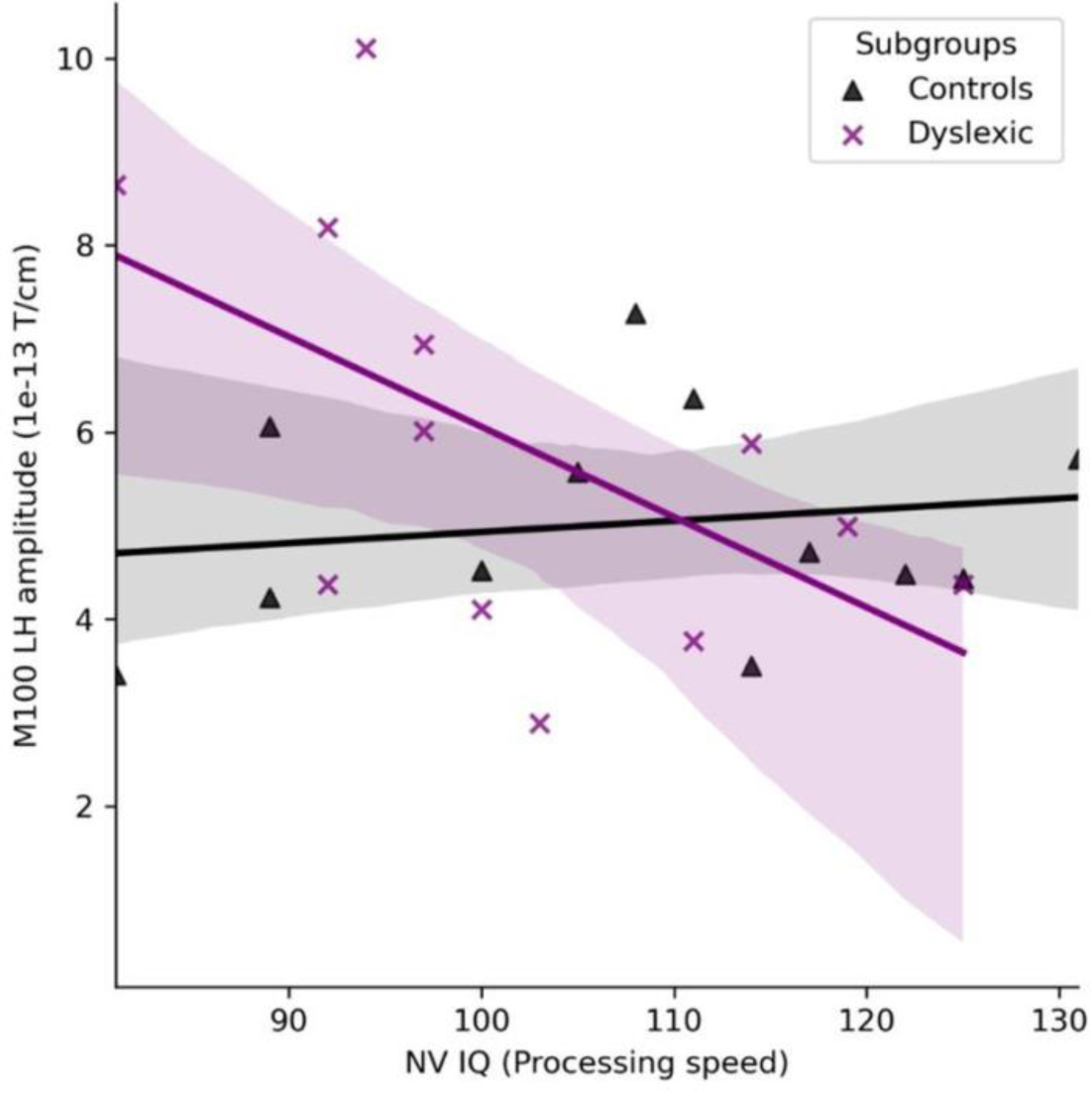
Scatter plot and correlations between NV IQ and the M100 left hemisphere (LH) amplitude, split by the controls/dyslexic grouping.

To explore the implications of these differences in the M100 left hemisphere amplitudes (for the dyslexic group) and right hemisphere latencies (for controls) on reading performance, we checked correlations between M100 measures and the PROLEC main score (see Supplementary Table S2). For dyslexic readers, the M100 LH amplitude was significantly correlated with the PROLEC main score, r = −0.619, p = 0.042. No other correlation, for both subgroups (controls and dyslexic), was significant.

### M200

The M200 latency had no significant main effects of Hemispheres, F(1, 20) = 0.312, p = 0.583, Controls/Dyslexic, F(1, 20) = 0.029, p = 0.866, and NV IQ, F(1, 20) = 1.125, p = 0.301. The only significant effect was the significant interaction of Hemispheres*NV IQ, F(1, 20) = 4.620, p = 0.044, but post hoc analysis did not reveal any significant differences. All the other interactions of Hemispheres*Controls/Dyslexic, F(1, 20) = 0.151, p = 0.702, Controls/Dyslexic*NV IQ, F(1, 20) = 0.075, p = 0.787, and the 3-way interaction Hemispheres*Controls/Dyslexic*NV IQ, F(1, 20) = 2.382, p = 0.138, were all not significant.

ANOVAs on the M200 amplitude also did not reveal any significant effects. The main effects of Hemispheres, F(1, 20) < 0.001, p = 0.988, Controls/Dyslexic, F(1, 20)=1.838, p=0.190, and NV IQ, F(1, 20) = 0.066, p = 0.800, were not significant. The interactions of Hemispheres*Controls/Dyslexic, F(1, 20) = 1.088, p = 0.309, Hemispheres*NV IQ, F(1, 20) = 4.090, p = 0.057, Controls/Dyslexic*NV IQ, F(1, 20) = 0.211, p = 0.651, and the 3-way interaction Hemispheres*Controls/Dyslexic*NV IQ, F(1, 20) = 0.946, p = 0.342, were also not significant.

### M400

The only significant effects recorded on the M400 component were the Hemisphere effect on latency, F(1, 20) = 7.875, p = 0.011, and amplitude, F(1, 20) = 6.417, p = 0.020. The latency of the M400 component was earlier for the right hemisphere compared to the left, while the amplitude was bigger for left hemisphere compared to the right. The main effects of Controls/Dyslexic, F(1, 20) = 0.023, p = 0.882, and NV IQ, F(1, 20) = 2.253, p = 0.149, were not significant. The interactions of Hemispheres*Controls/Dyslexic, F(1, 20) = 0.233, p = 0.634, Hemispheres*NV IQ, F(1, 20) = 0.392, p = 0.538, Controls/Dyslexic*NV IQ, F(1, 20) = 0.212, p = 0.650, in addition to the three-way interaction Hemispheres*Controls/Dyslexic*NV IQ, F(1, 20) = 3.388, p = 0.081, were also not significant. Similarly for the M400 amplitudes, the main effects of Controls/Dyslexic, F(1, 20) = 0.003, p = 0.960, and NV IQ, F(1, 20) = 0.458, p = 0.506, were not significant. Finally, the interactions of Hemispheres*Controls/Dyslexic, F(1, 20) = 0.059, p = 0.811, Hemispheres*NV IQ, F(1, 20) = 1.712, p = 0.206, Controls/Dyslexic*NV IQ, F(1, 20) = 0.626, p = 0.438, in addition to the three-way interaction Hemispheres*Controls/Dyslexic*NV IQ, F(1, 20) = 0.112, p = 0.741, were also not significant.

Finally, we note a significant effect of the Hemispheres*NV IQ interaction on the M200 latency, and a significant effect of hemisphere on the M400 latency and amplitude across the whole sample. However, post hoc analysis did not reveal any significant differences when corrected for multiple testing, suggesting these were driven by marginal effects.

## Discussion

The goal of this study was to gain a better understanding of how domain-general non-verbal cognitive skills captured by NV IQ differences interact with the auditory word processing in typical and dyslexic populations. We focused on the auditory word processing for two key reasons: (a) prior to and apart from reading deficits, dyslexia is linked to atypical auditory speech signal analysis hypothesized to contribute to reading deficits downstream; (b) developmental literature suggests that non-verbal skills affect aspects of the early stages of auditory word analysis. Therefore, we hypothesized that if effects of NV IQ are relevant for dyslexia, they are most likely to be present at the level of the cortical responses to auditory words. To test this, we used MEG data and ERF responses to understand whether different stages of auditory word processing are influenced by NV IQ. These distinct levels of auditory word processing were inspected through the M100, M200 and M400 ERF components elicited by listening to single words. To the best of our knowledge, this is the first study investigating NV IQ impact on typical adults and dyslexic readers in single word processing. The findings show neurophysiological evidence for NV IQ capacity being linked to the sensory processing of auditory stimuli as reflected by the M100 component.

The analysis contrasting the effect of NV IQ on controls and dyslexic participants revealed that the effect on the M100 latency and amplitude were different between the two groups. In auditory word paradigms, the M100 component is linked to early acoustic processing of spoken words (Näätänen & Picton, 1987). Our results showed that NV IQ is negatively correlated with the M100 latency in the RH for controls but not for the dyslexic group. Controls with lower NV IQ had delayed latencies compared to controls with higher NV IQ. This novel finding is consistent with previous literature showing that a shorter latency of the 100ms component is closely linked to mental ability and faster (and arguably better) processing in both verbal and non-verbal tasks (Pawlowski et al., 2019; Spironelli & Angrilli, 2009; Xue et al., 2017). The 100ms component has also been associated with attention allocation and working memory operation (Coull, 1998; Golob & Starr, 2004; Hillyard, et al., 1973). Faster allocation of these attention and working memory resources for auditory processing of language might be the reason behind the earlier RH M100 latency for controls with higher NV IQ (as measured with PSI -Processing Speed Index).

NV IQ was not the only possible contributor to this effect on the M100 latency. The speed of various cognitive processes declines with age (Schaie, 2013; Staff et al., 2014; Stuart-Hamilton, 2006) and this latter has been found to modulate latencies of the ERP components associated with early auditory responses (De Sanctis et al., 2008; Gajewski et al., 2018; Schapkin et al., 2014). To exclude this possibility, we both controlled for age within the dyslexic and control groups and conducted a supplementary regression analysis (Supplementary materials) that revealed that NV IQ is a significant predictor of the N100 latency in the RH over and above age and verbal IQ. While the discovered effect of NV IQ on the M100 latency is theoretically novel, it also has methodological implications for future studies. Many studies on language tend to control for age and verbal IQ; however, non-verbal IQ is rarely considered as an important factor. Our results highlight the importance of taking in consideration NV IQ in language related research, especially for studies exploring early stages of auditory processing.

The NV IQ effect on the M100 latency in controls was only present in the right hemisphere while no effect was recorded in the left hemisphere. Previous brain imaging studies have shown strong right hemisphere involvement in the processing of nonlinguistic perceptual features of verbal stimuli (Abrams et al., 2008; Belin et al., 2004; Hickok & Poeppel, 2007; Lattner et al., 2004). Furthermore, during auditory word processing, the left hemispheric dominance has primarily been shown for higher-order linguistic processes like semantic and syntactic processing (Friederici et al., 2003; Indefrey et al., 2001; Poldrack et al., 1999; Zahn et al., 2000). Consistently, our results have shown clear left lateralization at the M400 component which has been associated with lexico-semantic processing (Kutas & Federmeier, 2000; Van Petten et al., 1999). Accordingly, since the M100 component reflects early-stage processing of auditory stimuli, the RH effects of NV IQ are consistent with this narrative. Moreover, the RH has been argued to be part of the flexible periphery of brain regions linked to the core language network which may suggest broader involvement in various domain-general functions that support language processing (Bassett et al., 2013; Chai et al., 2016).

The absence of this effect in the dyslexic group can be interpreted as the result of deficits in mechanisms involved in initial sensory processing and early selective attention. For dyslexic readers, deficits in the early auditory processing were linked to reduced synchronization of the RH cortical signal with the auditory speech envelope (Molinaro et al., 2016; Power et al. 2013). According to the auditory temporal sampling hypothesis (Goswami, 2011), abnormal neural entrainment to speech at the low frequency auditory bands of the right auditory cortex contributes to poor speech signal segmentation and subsequent phonological deficits (Lizarazu et al., 2015; Power et al. 2013). Dyslexic readers’ difficulties in tracking speech rhythms in the right hemisphere (RH) is thought to hinder the phonological segmentation of a word into its underlying phonological representations and linking these elements to their corresponding sound in the LH (Lizarazu et al., 2021). Concurrently, dyslexic subjects have been shown to have deficiencies in attention (Moores et al., 2003) and working memory (Sela et al., 2012). Our findings are complementary with this literature showing that unlike controls, dyslexic readers do not show a benefit of faster auditory word analysis in the RH linked to stronger NV IQ skills. The absence of this link also implies that stronger NV IQ skills do not play compensatory role in terms of improved auditory processing speed in the dyslexic group.

While there was no NV IQ effect in latency for dyslexic readers, they displayed a significant effect of NV IQ on LH amplitudes. Dyslexic participants with lower NV IQ had larger LH M100 amplitudes compared to those with higher NV IQ. One interpretation of this effect is that lower NV IQ dyslexic readers need more cognitive resources for the early-stage auditory word processing in the LH. Similar hyper-activation in the LH (notably IFG) during auditory (Corina et al., 2001) and text processing (Morken et al; 2014) has been previously reported in dyslexia and linked to ‘inefficient processing’ strategies during reading. Here we show that hyperactivation in auditory word processing can further be linked to lower non-verbal skills. Differences between the dyslexic and control groups in the effect of NV IQ on latencies and amplitudes were only recorded for the M100 component while no significant differences were found between the two groups in the M200 and M400 components. This indicates that dyslexic readers exhibit selective processing anomalies at an earlier sensory level of auditory word analysis, with later-stage linguistic processing at the lexical and semantic level not affected by the non-verbal skills.

To explore the potential link between higher LH M100 amplitude found in the lower NV IQ dyslexic readers and their reading proficiency, we correlated LH M100 amplitudes with their PROLEC (main) scores. We found a mild negative correlation, r = −0.619, p = 0.042, suggesting that high LH M100 amplitudes are linked to worse reading skills in dyslexia. This supports our interpretation that the greater LH M100 amplitude likely represents less efficient (more resource-intensive) word-level auditory processing in dyslexia, further contributing to less efficient reading abilities. Unlike the latency results, the LH M100 amplitude findings are more in line with the compensatory view: in contrast to the lower NV IQ dyslexic group, dyslexic readers with higher NV IQ showed LH M100 amplitudes similar to those of the typical readers. In other words, if indeed non-verbal domain-general resources, especially those linked to the processing speed, can be harnessed to compensate for language-deficient skills, then the elevated LH M100 amplitude in the dyslexic participants with lower NV IQ could indicate a scarcity of such non-verbal cognitive resources available for efficient compensation, thus resulting in less effective control-like responses to auditory words. At the same time, we found no link between any of the neural measures linked to early-stage auditory word processing (LH/RH M100 amplitude or latency) in controls and their reading scores (see Supplementary Table S2) showing that early responses to auditory words are linked to reading only in the dyslexic group.

The analysis conducted on the M200 and M400 components did not reveal any significant NV IQ effects on their latencies or amplitudes for either the control or dyslexic groups. The only significant effect recorded, was the hemisphere lateralization effect on the M400. There was stronger activation in the left hemisphere compared to the right hemisphere indicating a left hemisphere lateralization during this stage of word processing. While the M100 reflects more sensory and perceptual processing, the M400 is arguably specialized for linguistic and lexico-semantic analysis (Zahn et al., 2000), and therefore is less affected by the domain-general skills.

## Conclusion

The present study explored the link between NV IQ and the neural signatures of auditory processing of single words in typical and dyslexic readers. Our findings provide novel evidence showing that non-verbal domain-general skills affects early stages of auditory word processing. Critically these effects were different for both groups with controls showing strong impact of NV IQ on the latency of the RH auditory word processing, and dyslexic readers on the amplitude of the early LH responses. These findings show that in addition to deficits in auditory processing, dyslexic readers also have atypical impact of domain-general cognitive skills on these processes. Future studies should actively assess the role of domain-general cognition in language processing, instead of controlling for the domain-general skills between populations.

## Acknowledgments

Authors would like to thank Manex Lete for his help with data collection.

## Supplementary Information

### Hierarchical regression

The effect of processing speed on the M100 right hemisphere latency for controls was further examined with a hierarchical regression. This regression is conducted in order to verify that the effect of NV IQ (as measured with PSI) is not already accounted for by age, verbal IQ (as measured with VCI) and the other IQ components (WMI & PRI). As such, the data met the assumptions of linearity, homoscedasticity, Breusch-Pagan test: Lagrange multiplier statistic = 5.564, p = 0.350, normal distribution of residuals, independence of errors, Durbin-Watson statistic = 1.486, p = 0.339, and collinearity (Age: Tolerance = 0.247, VIF = 4.053; VCI: Tolerance = 0.399, VIF = 2.508; WMI: Tolerance = 0.400, VIF = 2.500; PRI: Tolerance = 0.565, VIF = 1.770; PSI: Tolerance = 0.649, VIF = 1.540). Accordingly, the results of the regression indicate that NV IQ (measured with PSI) explained 29 %, R2 Change = 0.290, p =0.003, of the variance above the variance already explained by age, VCI, PRI and WMI. The regression model was a significant predictor of the right hemisphere latency for the M100 component, F(5, 11) = 15.869, p = 0.002, while PSI contributed significantly to the model, B = −5.807e −4, p = 0.003.

**Table. S1.** Correlations between PROLEC-SE-R main score and IQ indices for both the control (blue cells) and dyslexic (green cells) subgroups. Significant correlations are highlighted in bold.

|  | PSI | VCI | PRI | WMI | PROLEC-SE-R |
| --- | --- | --- | --- | --- | --- |
| **PSI** | - | r = 0.145,<br>p = 0.654 | r = 0.494,<br>p = 0.103 | r = 0.347,<br>p = 0.269 | r = 0.574,<br>p = 0.065 |
| **VCI** | r = -0.004,<br>p = 0.991 | - | r = 0.469,<br>p = 0.124 | **r = 0.642,**<br>**p = 0.024** | **r = 0.612,**<br>**p = 0.045** |
| **PRI** | r = -0.161,<br>p = 0.617 | r = 0.347,<br>p = 0.269 | - | r = 0.384,<br>p = 0.218 | r = 0.194,<br>p = 0.567 |
| **WMI** | r = -0.088,<br>p = 0.786 | r = 0.348,<br>p = 0.267 | **r = 0.634,**<br>**p = 0.027** | - | **r = 0.663,**<br>**p = 0.026** |
| **PROLEC-SE-R** | r = 0.555,<br>p = 0.061 | **r = 0.602,**<br>**p = 0.038** | r = 0.038,<br>p = 0.907 | r = 0.263,<br>p = 0.409 | - |

**Table. S2.**
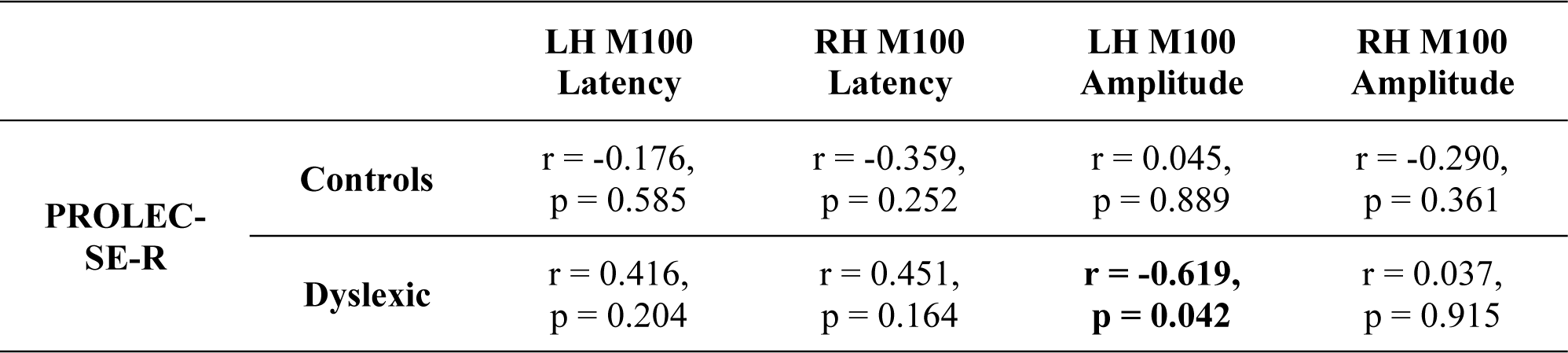
Correlations between PROLEC-SE-R main score and the M100 component (latency and amplitude for each hemisphere). Significant correlations are highlighted in bold.

